# Screening and identification of muscle-specific candidate genes via mouse microarray data analysis

**DOI:** 10.1101/2021.08.11.456020

**Authors:** Sayed Haidar Abbas Raza, Chengcheng Liang, Wang Guohua, Linsen Zan

**Affiliations:** College of Animal Science and Technology, Northwest A&F University, Yangling, Shaanxi, 712100, China; National Beef Cattle Improvement Center, Northwest A&F University, Yangling, Shaanxi, 712100, China

**Keywords:** Muscle development, Microarray analysis, Differential genes, Bioinformatics

## Abstract

Muscle tissue is involved with every stage of life activities and has roles in biological processes. For example, the blood circulation system needs the heart muscle to transport blood to all parts, and the movement cannot be separated from the participation of skeletal muscle. However, the process of muscle development and the regulatory mechanisms of muscle development are not clear at present. In this study, we used bioinformatics techniques to identify differentially expressed genes specifically expressed in multiple muscle tissues of mice as potential candidate genes for studying the regulatory mechanisms of muscle development. Mouse tissue microarray data from 17 tissue samples was selected from the GEO database for analysis. Muscle tissue as the treatment group, and the other 16 tissues as the control group. Genes expressed in the muscle tissue were different to those in the other 16 tissues and identified 272 differential genes with highly specific expression in muscle tissue, including 260 up-regulated genes and 12 down regulated genes. is the genes were associated with the myofibril, contractile fibers, and sarcomere, cytoskeletal protein binding, and actin binding. KEGG pathway analysis showed that the differentially expressed genes in muscle tissue were mainly concentrated in pathways for AMPK signaling, cGMP PKG signaling calcium signaling, glycolysis, and, arginine and proline metabolism. A PPI protein interaction network was constructed for the selected differential genes, and the MCODE module used for modular analysis. Five modules with Score > 3.0 are selected. Then the Cytoscape software was used to analyze the tissue specificity of differential genes, and the genes with high degree scores collected, and some common genes selected for quantitative PCR verification. The conclusion is that we have screened the differentially expressed gene set specific to mouse muscle to provide potential candidate genes for the study of the important mechanisms of muscle development.

## 1. Introduction

Muscles represent a crucial group of soft tissues derived from the mesoderm that are primarily responsible for locomotion, and movement in all animals. The World Health Organization estimates musculoskeletal disorders cause the highest proportion of disabilities worldwide, affecting approximately 1.7 billion people (Cieza et al., 2020). Therefore, there is significant interest in characterizing the genetics that underpin muscular development, and any associated pathophysiology.

There are three major group of muscles i.e. skeletal, myocardium and smooth muscles. Skeletal muscles weigh about 40% of adult weight in humans, and represent the main subgroup of muscles that allow for locomotion in conjunction with the skeletal system. Apart from locomotion, skeletal muscles also have other important functions e.g. heat production, support and protection of other soft tissues, and participation in metabolic homeostasis (Cohen et al., 2015; Wang et al., 2019). Diseases that affect primary skeletal muscles or the neuromuscular junction frequently manifest in the form of pathological muscle weakness or reduced skeletal muscle mass, which weakens the body’s ability to respond to stress and chronic diseases (Frontera and Ochala, 2015). Moreover, amino acids released from muscles help maintain blood sugar levels during starvation. Therefore, diseases affecting skeletal muscles can result in wide ranging pathologies, and represent a key cause of morbidity and disability in human populations.

Skeletal muscles are multinucleated, and develop via the fusion of myogenic progenitor cells called myoblasts, into muscle fibers called myotubes, via a complex process known as myogenesis (Buckingham and Rigby, 2014; Wang and Rudnicki, 2012). Several genes are known to play a crucial role either during myogenesis, or subsequently, in ensuring normal muscle physiology (Horak et al., 2016). Some of the main genes involved in muscular development include transcription factors *MYOD1* (myogenic differentiation 1), *MYF5* (myogenic factor 5), *MYOG* (myogenin) and *MRF* (myogenic regulatory factor), *MYF6* (herculin), *PAX3* (paired box 3), *PAX7* (paired box 7) and *MEF2* (myocyte enhancer factor 2) family (Brand-Saberi, 2005). *MYOD1* and *MYF5* are involved in the early phases of skeletal muscle development by promoting the proliferation and differentiation of myogenic progenitor cells into myoblasts, while *MYOG* plays an important role in the latter phases of myogenesis that involve fusion of myoblasts into myotubes. The precise function of *MYF6* remains unknown, though it is thought to regulate myogenesis, and is exclusively expressed in skeletal muscles (Ito et al., 2012).

Apart from these widely known genes, several other genes that influence either skeletal muscle development or physiology remain unidentified and/or uncharacterized.

High-throughput gene chip technologies that provide large-scale gene expression data by measuring transcript abundance in various tissues or cells (Barrett et al., 2005), can be leveraged in combination with online gene expression databases (e.g. NCBI’s GEO database) to identify such genes. Moreover, given that human populations are genetically heterogenous, inbred animals models can be very useful in identifying and characterizing key genes associated with muscle development and disease.

Therefore, the overall aim of the present study, was to identify genes that are differentially expressed in skeletal muscles of 10-12 week old C57BL/6 mice, by comparing skeletal muscle expression profiles against 16 non-muscle tissues. Genes identified as differentially expressed in muscles, were subsequently subjected to bioinformatic analyses including process and pathway enrichment analysis, protein-protein interaction (PPI) network construction and molecular compounding. Finally, the genes with partial height difference multiples were selected for validation via qPCR.

## 2. Materials and methods

### 2.1 Ethics statement

All procedures were approved by the Experimental Animal Center of Xi’an Jiaotong University. Animal care and use protocols (EACXU 172) were approved by the Institutional Animal Care and Use Committee of Xi’an Jiaotong University. All animal experiments were performed in adherence with the NIH Guidelines on the Use of Laboratory Animals.

### 2.2 Microarray data

The microarray data was downloaded from NCBI’s GEO (Gene Expression Omnibus) database (GEO accession number GSE9954). The downloaded dataset contained microarray expression data from 70 samples that collectively represent 22 tissues (including muscles). The microarray dataset was derived from 10-12 week old male C57BL/6 mice using the Affymetrix Mouse Genome 430 2.0 Array platform (GPL1261). After euthanasia, multiple organs and tissues were taken for microarray analysis (Thorrez et al., 2008). In this study, microarray data from 17 out of the 22 available tissues was selected. The selected tissues included muscles, adipose tissue, adrenal gland, bone marrow, brain, eye, heart, kidney, liver, lung, pituitary gland, placenta, salivary gland, seminal vesicle, small intestine, spleen, testis, and thymus.

Differential gene expression analysis was performed on the downloaded microarray data using the R project for statistical computing (version 3.5.2; https://www.r-project.org/) packages ‘limma’ (http://www.bioconductor.org/packages/3.5/bioc/html/limma.html) (Troyanskaya et al., 2001), and ‘impute’ (http://www.bioconductor.org/packages/2.7/bioc/html/impute.html) (Ritchie et al., 2015). Screening for differentially expressed genes (DEGs) was performed by comparing mouse muscle expression profiles against the remaining 16 tissues using a p-value threshold of <0.05, and log2(fold change) threshold of ≥2.

### 2.3 Process and pathway enrichment analyses

Genes identified as differentially expressed in the initial screening, were subjected to Gene Ontology (GO) analysis via the Database for Annotation, Visualization and Integrated Discovery (DAVID) (https://david.ncifcrf.gov/home.jsp) using the *Mus musculus* genome annotation as background. Three aspects of the GO database were targeted in the GO enrichment analyses i.e. cellular component (CC), molecular function (MF), and biological process (BP). Similarly, KEGG pathway enrichment analysis was performed using DAVID and KOBAS (KEGG Orthology-Based Annotation System - http://kobas.cbi.pku.edu.cn/). The R package ‘ggplot2’ (version: 3.1.0; http://ggplot2.tidyverse.org) was used for data visualization.

### 2.4 PPI network and module analysis

Protein-protein interaction network and module analysis was performed using online tools and the String database (https://string-db.org/). Genes identified as differentially expressed in muscles were used to construct a PPI network map (Franceschini et al., 2012), and the MCODE (Molecular Complex Detection) plug-in of Cytoscape software (version: 3.6.0; Java version: 1.8.0_201) was subsequently used to identify interconnected clusters within the PPI network using a node cutoff score of >3.0. The top 30 proteins with the highest number of degrees (i.e. edges) were represented in the form of a bar graph (Shannon et al., 2003); and network modules identified via MCODE (score >3.0) were also represented diagrammatically (Bader and Hogue, 2003).

### 2.5 Animals and tissues collection

The animals used in this study were obtained from the Experimental Animal Center of the Medical College of Xi’an Jiaotong University. As per approved animal use protocols 12-week-old female C57B/L mice were euthanized with 5% chloral hydrate, and tissue samples (heart, liver, spleen, lung, kidney, muscles and adipose) were subsequently collected surgically. All surgical instruments used in the experiment were put into 0.1% DEPC solution overnight, and then autoclaved and dried for use. Collected tissue samples were rinsed in pre-chilled Phosphate Buffer Saline (PBS), put into RNase-free centrifuge tubes, and immediately snap frozen in liquid nitrogen. Total RNA was extracted from these tissue samples after transportation to the laboratory.

### 2.6 Total RNA extraction and cDNA

Frozen tissue samples were homogenized prior to RNA extraction using enzyme-free centrifuge tubes containing Trizol (TakaraBio, Dalian, China), as per manufacturers instructions. The concentration of the extracted total RNA was determined via nanodrop quantification. Finally, extracted RNA samples were reverse transcribed into cDNA using a Prime Script RT Reagent Kit (TakaraBio, Dalian, China) for subsequent quantitative PCR.

### 2.7 Primer information

Intron spanning primers were designed using Primer Premier ver. 5.0 (PREMIER Biosoft, http://www.premierbiosoft.com/). The primer sequences, as well as annealing temperatures are described in Table 5.

### 2.8 qRT-PCR

The CFX-96 (BIO-RAD, US) was used to carry out real-time fluorescence quantitative polymerase chain reaction (qRT-PCR) using a commercially available kit (TB Green Premix Ex Taq II, Tli RNaseH Plus, TakaraBio, Dalian, China). Three reference genes (*18s rRNA, GAPDH* and *β-actin*) were tested as internal controls via homogeneity checks. Subsequently, the geometric mean values of *18s rRNA* and *β-actin* was decided to be used as internal reference for qRT-PCR. The final raw data was analyzed via the delta-delta Ct (2^-ΔΔCT^) calculation method (Livak and Schmittgen, 2001), and graphical analysis of data was performed via GraphPad Prism 6.0 (https://www.graphpad.com/scientific-software/prism/).

## 3. Results

### 3.1 Differentially expressed genes (DEGs) analysis

Microarray data was normalized prior to differentially expressed genes (DEGs) analysis. The gene expression profiles prior to normalization, and after normalization are presented in Figures 1A and 1B respectively. A volcano plot showing differentially expressed genes (DEGs) that were upregulated or downregulated when contrasting muscle gene expression profiles against a combination of the remaining 17 tissues, is presented in Figure 1C.

**Figure 1.**
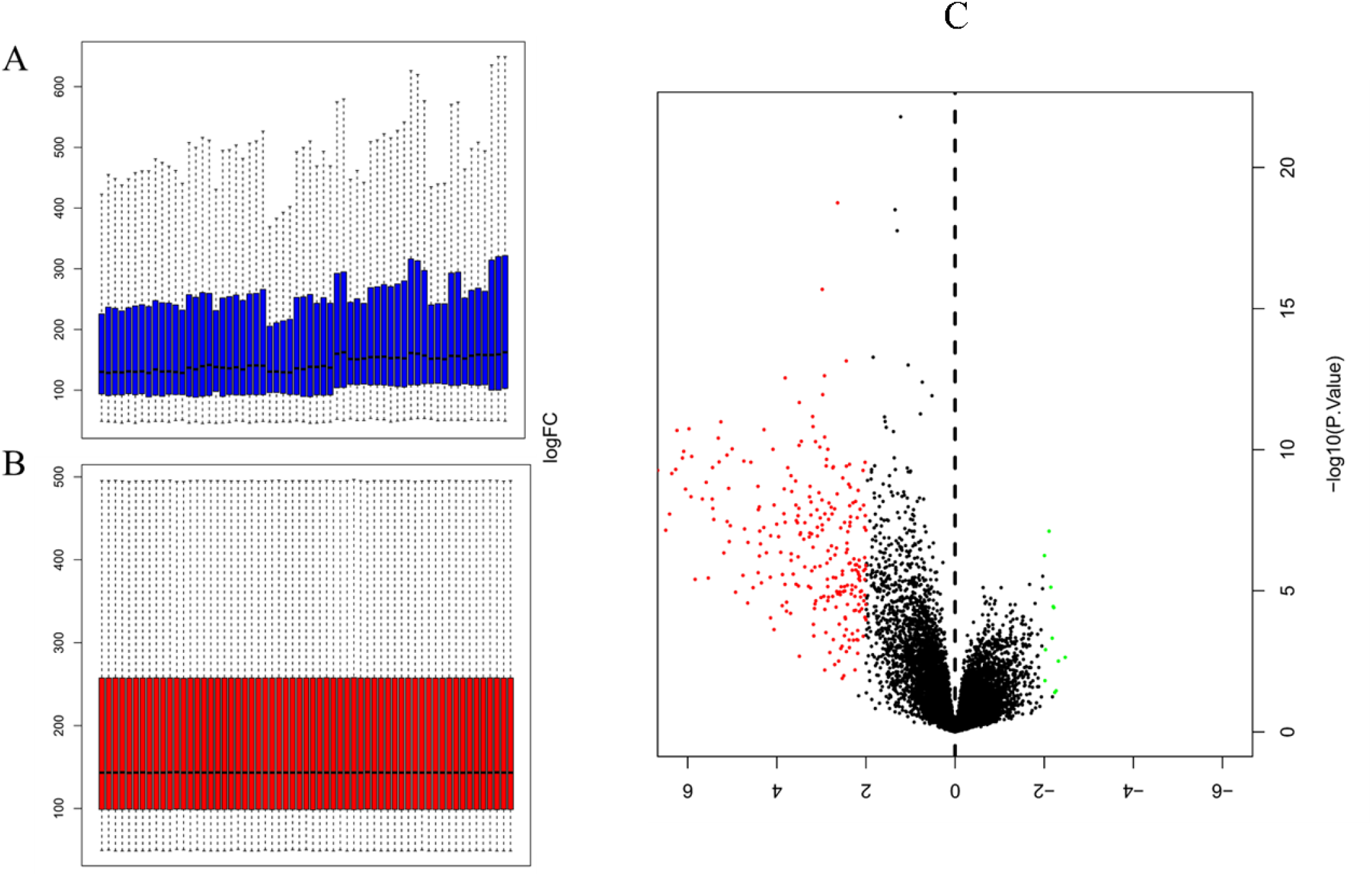
Normalization of microarray expression data, and volcano plot of DEGs (A) Gene expression profile prior to normalization (B) and after normalization was carried out using Limma package in R; (C) Volcano plot of DEGs (P<0.05 and log2|FC|≥2) identified by contrasting the expression profiles in muscles against the combined expression profiles of 17 other tissues. The y-axis represents the log2|FC|, and the x-axis displays the statistical significance of the differences. Black dots represent genes that were not found to be differentially expressed. Red dots represent genes that were significantly upregulated, and green dots represent genes that were significantly downregulated.

Genes that were significantly up or downregulated (p value <0.05) in muscles, with a log2(fold change) ≥2, were identified as differentially expressed genes (DEGs). Gene expression profile of muscles was first contrasted against each of the 17 control tissues used in this study. The number of differentially expressed genes (DEGs) identified in each of these individual contrasts are noted in Table 1. Comparing, gene expression in muscle to the expression profiles of all 17 control tissues combined, allowed for the identification of 260 DEGs that were upregulated, and 12 differentially expressed genes (DEGs) that were downregulated in muscle. Some of the DEGs that were found to be upregulated include myocyte differentiation markers myosin light chain 1 (*Myl1*), myosin heavy chain 4 (*Myh4*), myosin heavy chain 2 (*Myh2*), and inositol protein (*Myot*). Myosin heavy chain 1 (*Myh1*) was found to have the highest log2 (fold change) of 6.675, which is indicative of a more than 100-fold greater expression in muscles relative to the combination of the 17 control tissues used in this study. Amongst the genes that were downregulated, Cytochrome C Oxidase Subunit 6A1 (*Cox6a1*) was found to have the lowest log2 (FC) of -2.472, which is equivalent to an approximately 5-fold reduction in gene expression. A complete list of the top 20 upregulated DEGs, and all of the 12 downregulated DEGs, is presented in Table 2.

**Table 1:**
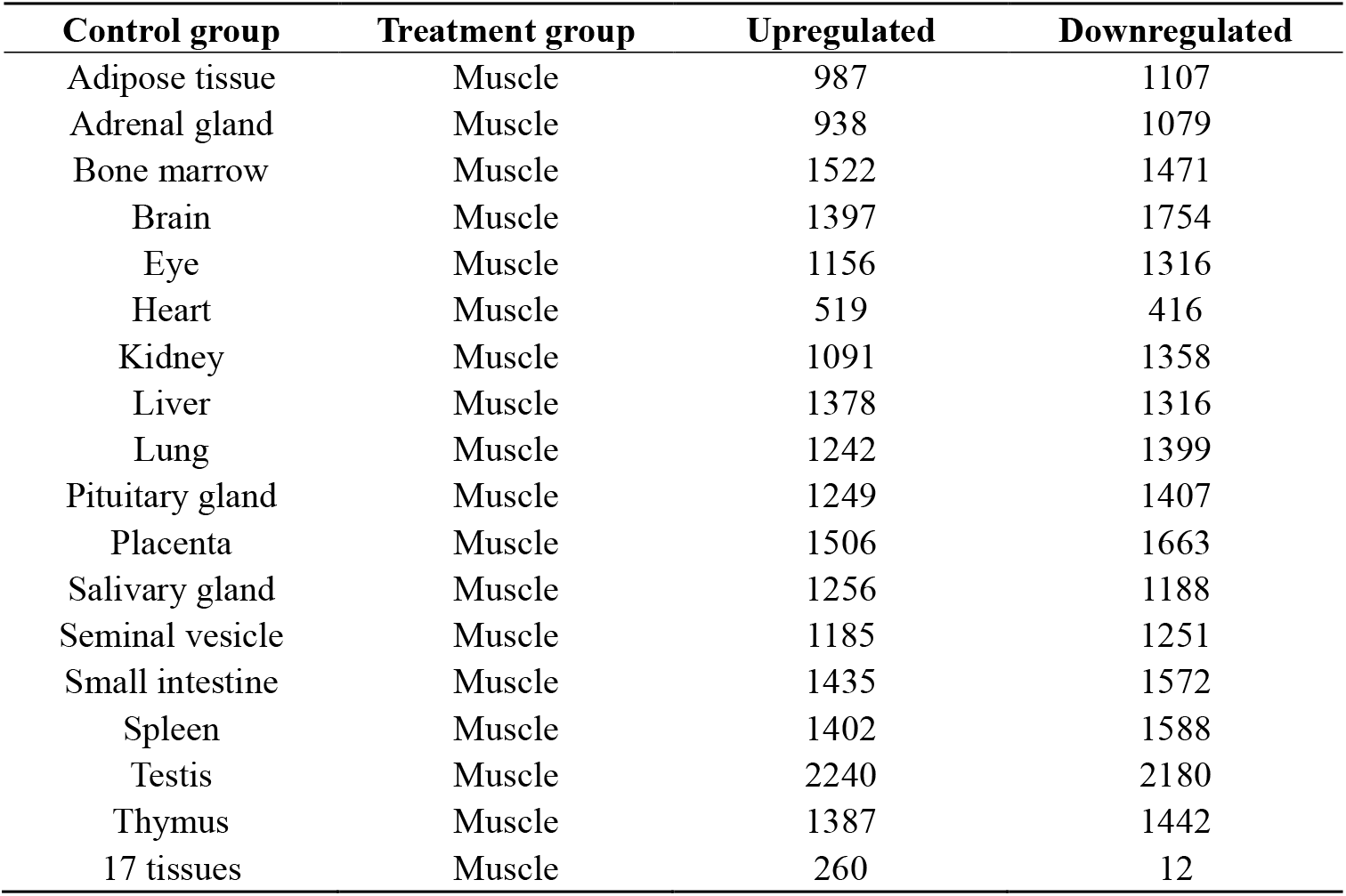
The number of upregulated and downregulated DEGs identified by contrasting muscle expression profiles against the expression profiles of different tissues.

**Table 2:**
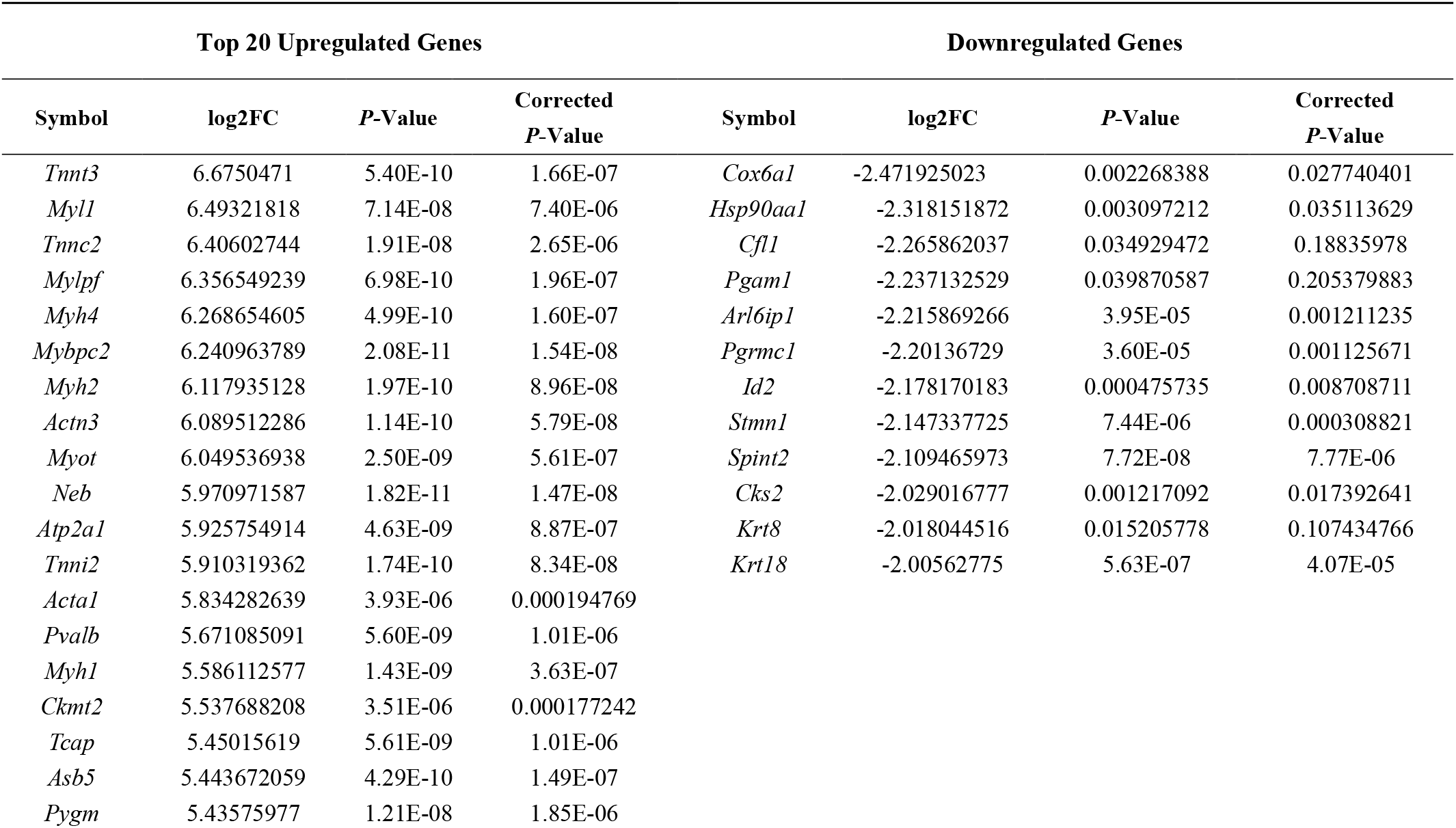
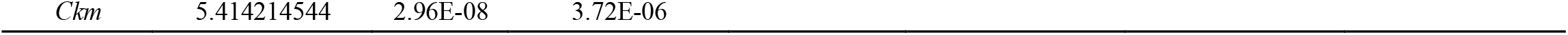
Fold change and statistical significance the top 20 upregulated, all 12 downregulated DEGs when comparing muscle expression profile against the combined expression profile of remaining 17 tissues.

| Top 20 Upregulated Genes |  |  |  | Downregulated Genes |  |  |  |
| --- | --- | --- | --- | --- | --- | --- | --- |
| Symbol | log2FC | P-Value | Corrected P-Value | Symbol | log2FC | P-Value | Corrected P-Value |
| *Tnnt3* | 6.6750471 | 5.40E-10 | 1.66E-07 | *Cox6a1* | -2.471925023 | 0.002268388 | 0.027740401 |
| *Myl1* | 6.49321818 | 7.14E-08 | 7.40E-06 | *Hsp90aa1* | -2.318151872 | 0.003097212 | 0.035113629 |
| *Tnnc2* | 6.40602744 | 1.91E-08 | 2.65E-06 | *Cfl1* | -2.265862037 | 0.034929472 | 0.18835978 |
| *Mylpf* | 6.356549239 | 6.98E-10 | 1.96E-07 | *Pgam1* | -2.237132529 | 0.039870587 | 0.205379883 |
| *Myh4* | 6.268654605 | 4.99E-10 | 1.60E-07 | *Arl6ip1* | -2.215869266 | 3.95E-05 | 0.001211235 |
| *Mybpc2* | 6.240963789 | 2.08E-11 | 1.54E-08 | *Pgrmc1* | -2.20136729 | 3.60E-05 | 0.001125671 |
| *Myh2* | 6.117935128 | 1.97E-10 | 8.96E-08 | *Id2* | -2.178170183 | 0.000475735 | 0.008708711 |
| *Actn3* | 6.089512286 | 1.14E-10 | 5.79E-08 | *Stmn1* | -2.147337725 | 7.44E-06 | 0.000308821 |
| *Myot* | 6.049536938 | 2.50E-09 | 5.61E-07 | *Spint2* | -2.109465973 | 7.72E-08 | 7.77E-06 |
| *Neb* | 5.970971587 | 1.82E-11 | 1.47E-08 | *Cks2* | -2.029016777 | 0.001217092 | 0.017392641 |
| *Atp2a1* | 5.925754914 | 4.63E-09 | 8.87E-07 | *Krt8* | -2.018044516 | 0.015205778 | 0.107434766 |
| *Tnni2* | 5.910319362 | 1.74E-10 | 8.34E-08 | *Krt18* | -2.00562775 | 5.63E-07 | 4.07E-05 |
| *Acta1* | 5.834282639 | 3.93E-06 | 0.000194769 |  |  |  |  |
| *Pvalb* | 5.671085091 | 5.60E-09 | 1.01E-06 |  |  |  |  |
| *Myh1* | 5.586112577 | 1.43E-09 | 3.63E-07 |  |  |  |  |
| *Ckmt2* | 5.537688208 | 3.51E-06 | 0.000177242 |  |  |  |  |
| *Tcap* | 5.45015619 | 5.61E-09 | 1.01E-06 |  |  |  |  |
| *Asb5* | 5.443672059 | 4.29E-10 | 1.49E-07 |  |  |  |  |
| *Pygm* | 5.43575977 | 1.21E-08 | 1.85E-06 |  |  |  |  |

### 3.2 Process and pathway enrichment analysis

. Enrichment analysis targeting GO terms identified a total of 752 GO annotations that were significantly enriched (p-value <0.01) in differentially expressed genes DEGs identified within this study. Of the total 752 GO terms, 548 GO terms represented biological processes amongst which, the most significantly enriched processes included muscle system process, muscle structure development, myofibril assembly and muscle cell development. A further 103 GO terms representing cellular components were identified, of which the most significantly enriched GO terms including myofibrils, contractile fibers, sarcomeres and contractile fibers. Finally, a total of 101 GO terms representing molecular functions were identified, of which, the most significantly enriched GO terms included cytoskeletal protein binding, actin binding, and structural molecular functions. A summary of the top 10 GO terms identified in each of the three GO aspect categories is presented in Table 3. Top GO terms identified through enrichment analysis are also presented diagrammatically in Figure 2. Complete enrichment analysis results are presented in supplementary Table S2.

**Table 3:** GO (Gene Ontology) enrichment analysis of identified DEGs.

| Ontology | GO ID | GO Term | Count | P-Value | FDR |
| --- | --- | --- | --- | --- | --- |
| MF | GO:0008092 | cytoskeletal protein binding | 71 | 2.56E-36 | 1.68E-33 |
|  | GO:0003779 | actin binding | 45 | 3.55E-29 | 1.16E-26 |
|  | GO:0008307 | structural constituent of muscle | 13 | 1.00E-19 | 2.20E-17 |
|  | GO:0051015 | actin filament binding | 25 | 8.69E-19 | 1.42E-16 |
|  | GO:0005515 | protein binding | 191 | 1.36E-17 | 1.79E-15 |
|  | GO:0051371 | muscle alpha-actinin binding | 9 | 9.88E-15 | 1.08E-12 |
|  | GO:0042805 | actinin binding | 12 | 6.72E-14 | 6.30E-12 |
|  | GO:0051393 | alpha-actinin binding | 10 | 3.12E-12 | 2.56E-10 |
|  | GO:0031432 | titin binding | 7 | 1.17E-11 | 8.55E-10 |
|  | GO:0005523 | tropomyosin binding | 7 | 5.98E-11 | 3.92E-09 |
| BP | GO:0003012 | muscle system process | 63 | 1.40E-50 | 4.81E-47 |
|  | GO:0061061 | muscle structure development | 71 | 1.33E-47 | 2.30E-44 |
|  | GO:0030239 | myofibril assembly | 33 | 1.44E-46 | 1.66E-43 |
|  | GO:0055001 | muscle cell development | 45 | 1.81E-45 | 1.56E-42 |
|  | GO:0055002 | striated muscle cell development | 43 | 3.36E-44 | 2.32E-41 |
|  | GO:0006936 | muscle contraction | 51 | 6.66E-44 | 3.83E-41 |
|  | GO:0051146 | striated muscle cell differentiation | 48 | 1.24E-39 | 6.10E-37 |
|  | GO:0010927 | cellular component assembly involved in morphogenesis | 33 | 1.77E-38 | 7.61E-36 |
|  | GO:0042692 | muscle cell differentiation | 51 | 2.04E-38 | 7.83E-36 |
|  | GO:0007517 | muscle organ development | 49 | 2.17E-35 | 7.49E-33 |
| CC | GO:0030016 | myofibril | 78 | 4.81E-96 | 1.69E-93 |
|  | GO:0043292 | contractile fiber | 79 | 7.03E-96 | 1.69E-93 |
|  | GO:0030017 | sarcomere | 74 | 1.06E-92 | 1.51E-90 |
|  | GO:0044449 | contractile fiber part | 75 | 1.25E-92 | 1.51E-90 |
|  | GO:0031674 | I band | 51 | 5.63E-62 | 5.42E-60 |
|  | GO:0030018 | Z disc | 43 | 2.59E-50 | 2.08E-48 |
|  | GO:0099512 | supramolecular fiber | 83 | 1.01E-49 | 6.93E-48 |
|  | GO:0099081 | supramolecular polymer | 83 | 1.53E-49 | 8.90E-48 |
|  | GO:0099080 | supramolecular complex | 83 | 1.67E-49 | 8.90E-48 |
|  | GO:0005865 | striated muscle thin filament | 21 | 1.45E-35 | 6.96E-34 |

**Figure 2.**
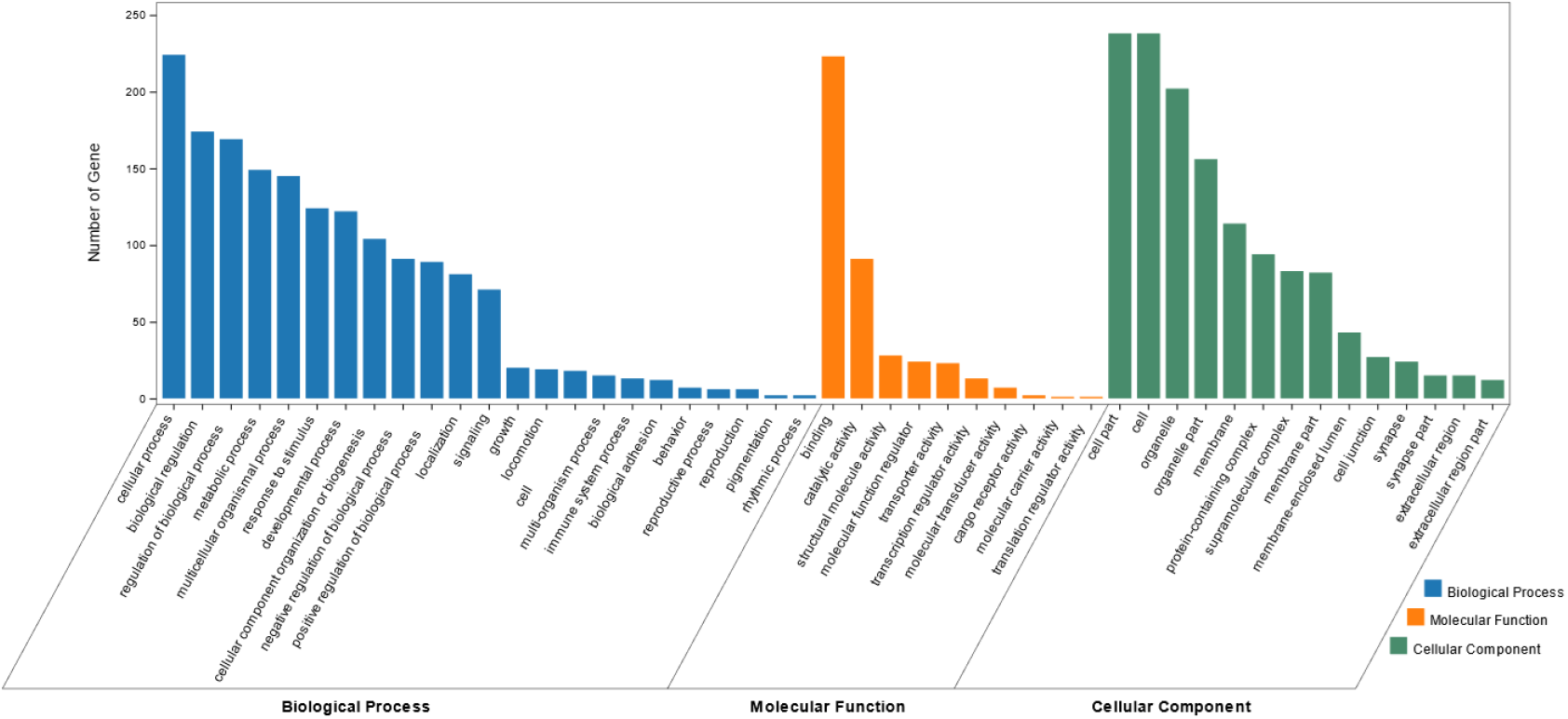
Bar graph of the most enriched GO terms of identified DEGs. The x-axis represent the top Biological Processes (BP), Cellular Components (CC) and Molecular Functions (MF) identified via GO enrichment analysis. The y-axis represents the number of DEGs identified within each GO term.

Functional enrichment analysis performed using DAVID identified 29 KEGG pathways significantly associated with muscle specific differentially expressed genes DEGs (P-value <0.01). The top 20 of these KEGG pathways are presented in the form of a bubble chart in Figure 3, which demonstrates that the identified differentially expressed genes DEGs are highly relevant in cardiac function and pathophysiology. Other KEGG pathways identified via DAVID analysis included AMPK, cGMP-PKG and calcium signaling pathways; Glycolysis / Gluconeogenesis, Carbon metabolism, Arginine and Proline metabolism etc. (Figure 3)

**Figure 3.**
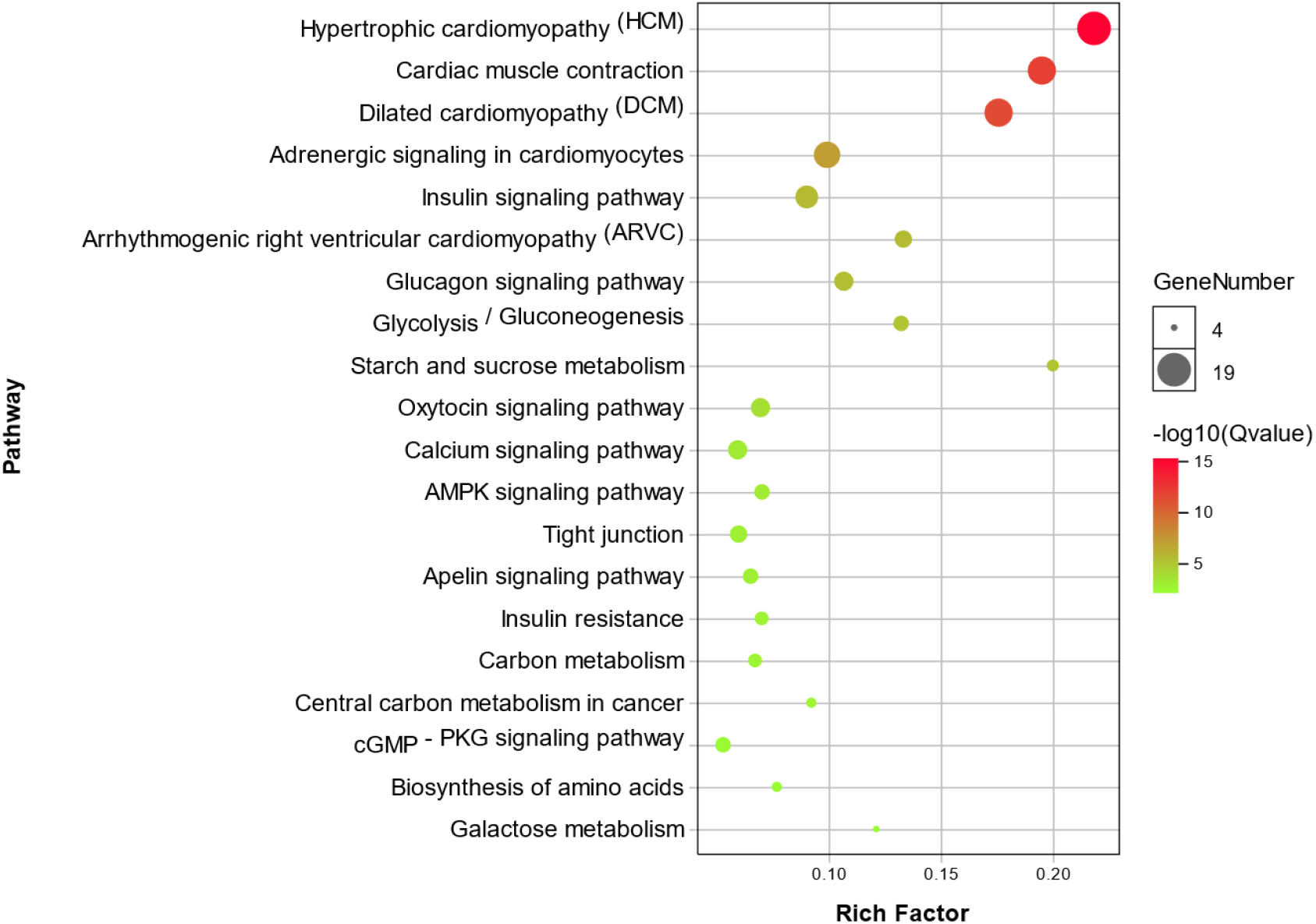
Bubble chart showing enrichment of DEGs in the top 20 KEGG pathways. The y-axis represents different KEGG pathways that were found to be enrichment in the identified DEGs. The x-axis represents the rich factor, which in turn represents the ratio of DEGs to the total number of genes in any given KEGG pathway. The size of each bubble represents the number of DEGs in a given KEGG pathway, and the color represents enrichment significance.

### 3.3 PPI network analysis and module screening

Network analysis focused on protein-protein interactions performed using the online STRING (https://string-db.org/) database, identified a total of 247 Nodes and 2,813 Edges (score>0.4). Results from the PPI network analysis are presented diagrammatically in Figure 4A, which shows upregulated DEGs in red, and downregulated DEGs in blue, with the color intensity corresponding to fold changes (darker colors reflecting higher fold changes). These results clearly indicate the presence of a large highly correlated network of upregulated muscle specific genes. Network analysis was further performed via Cytoscape software to compute the number of connections of each individual node (i.e. node degrees), and these results are presented in Table S4. The top 30 nodes with the highest number of connections (i.e. degrees), presented in Figure 4B, were comprised by Titin (*Ttn*, 103 degrees); Actinin Alpha 2 (*Actn2*, 86 degrees); Creatine Kinase, Mitochondrial 2 (*Ckmt2*, 83 degrees); LIM Domain Binding 3 (*Ldb3*, 83 degrees); Muscle Creatine Kinase (*Ckm*, 81 degrees); Obscurin, Cytoskeletal Calmodulin and Titin-Interacting RhoGEF (*Obscn*, 80 degrees); and Titin-Cap (*Tcap*, 80 degrees), in addition to many other muscle-specific DEGs.

**Figure 4.**
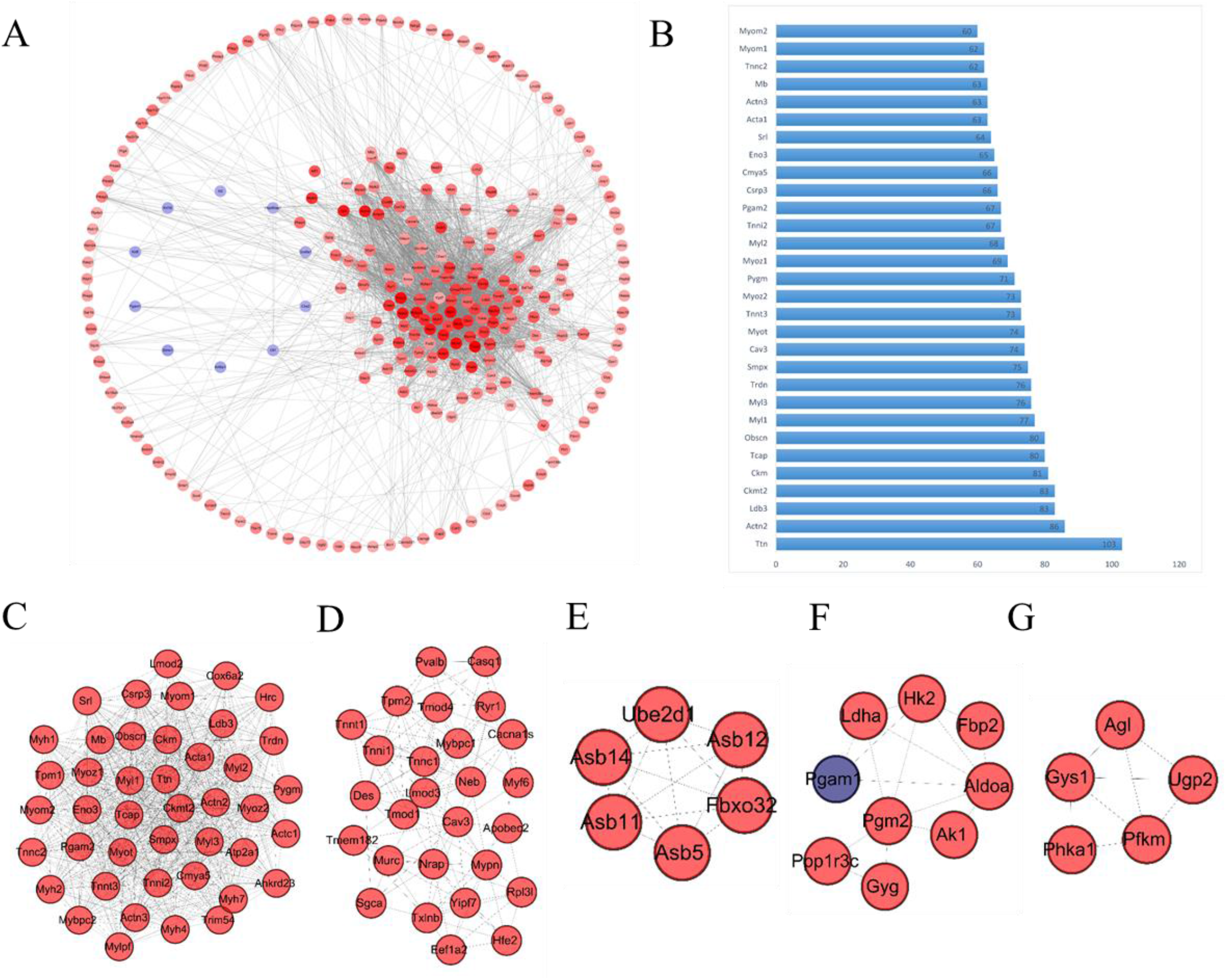
PPI network and modules (A) Overall PPI network constructed using the identified DEGs. Genes represented in red are upregulated, and those represented in blue are downregulated within the network. The color intensity represents fold change of gene expression, with higher color intensity representing a higher fold change in expression levels. (B) Bar graph representing the top 30 genes with the highest degrees (connections) within the PPI network; the y-axis represents each of the top 30 genes, and the x-axis represents the number of connections of each gene within the network (degrees). (C-G) Network modules identified via MCODE, including module 1 (score 36.905), module 2 (score 9.769), module 3 (score 6), module 4 (score 4) and module 5 (score 4). Again genes represented in red indicate upregulation, and genes represented in blue indicate downregulation.

Network module analysis performed via MCODE plug-in of Cytoscape, identified a further five modules (module score > 3.0), which are presented in Figures 4C-G. Module 1 (Figure 4C) has the highest MCODE score of 36.905, and included 43 interacting proteins (nodes) with 775 interactions (edges). The second module (Figure 4D) was considerably smaller with an MCODE score of 9.769, including 27 nodes and 127 edges. As evident in Figures 4C-G, most of the DEGs represented in these modules were upregulated (Red nodes indicate upregulated nodes, and blue indicates downregulated nodes).

### 3.4 qRT -PCR validation of identified DEGs

Differentially expressed genes DEGs that were identified in this study were also annotated for tissue specific expression using the online DAVID (https://david.ncifcrf.gov/) database, and this identified 55 genes with tissue specific expression in skeletal muscles (Table 4). A Venn diagram constructed to compare these 55 genes against the list of top 30 genes identified via Cytoscape network analysis, identified 11 genes shared in common (Figure 5). These genes included: Actinin Alpha 2 (*ACTN2*), LIM Domain-Binding Protein 3 (*LDB3*), Small Muscle Protein X-Linked (*SMPX*), Caveolin 3 (*CAV3*), Troponin T3, Fast Skeletal Type (*TNNT3*), Myozenin 2 (*MYOZ2*), Glycogen Phosphorylase, Muscle Associated (*PYGM*), Cardiomyopathy Associated 5 (*CMYA5*), Enolase 3 (Beta, Muscle) (*ENO3*), Sarcalumenin (*SRL*) and Actinin Alpha 3 (*ACTN3*).

**Table 4:** Tissue specific expression annotations of differentially expressed genes DEGs identified via DAVID analysis.

| Tissues | Count | P-Value | Genes |
| --- | --- | --- | --- |
| Skeletal muscle | 55 | 2.15E-57 | *PRKAG3, PDLIM3, ANKRD2, RTN2, ART3, JSRP1, PVALB, SH3BGR, JPH1, RBFOX1, MYH1, MYH2, TNMD, LDB3, MYH4, ACTN2, MYH7, FBP2, ACTN3, CACNG1, TACC2, TNNT3, TNNT1, SGCG, CFL2, ITGB1BP2, RYR1, SMPX, SGCA, SGCB, CAV3, FHL1, PHKA1, SRL, TPM2, KCNA7, ENO3, DDIT4L, HRC, MYF6, MUSTN1, MYOZ2, TRIM63, TNNI1, SLC16A3, TUBA8, NEB, NRAP, MAPK12, PYGM, GYG, ATP2A1, CMYA5, SYNM, VLDLR* |
| Heart | 58 | 8.55E-19 | *APOBEC2, LDHA, TNNC1, PGAM2, ANKRD1, TXLNB, TTN, ART1, LMOD2, PPP1R14C, USP13, HSP90AA1, SLC25A4, CRYAB, LDB3, ACTN2, MYH7, GMPR, IRS1, TRDN, MURC, HSPB6, CFL2, HSPB8, HSPB7, SMPX, TCAP, MYL3, SMTNL2, ASB14, TPM1, ASB15, KCNA7, MYOM2, CKMT2, MLIP, POPDC3, HRC, LPL, ADSSL1, ACTC1, CACNA2D1, COX8B, ALPK3, YIPF7, PDK4, ATP1A2, TRIM63, CSRP3, CACNA1S, IDH3A, FSD2, ABCC9, PTP4A3, FABP3, COX6A2, KLHL30, VLDLR* |
| Muscle | 12 | 3.50E-11 | *RBFOX1, ADSSL1, SLC25A4, SLC2A4, PHKG1, CMYA5, HSPB7, KY, SYNPO2, MYOM1, MYOT, SNTA1* |
| Heart muscle | 5 | 2.21E-06 | *ART3, TRIM54, NRAP, XIRP1, MYL2* |
| Bone | 15 | 1.54E-05 | *PRKAG3, FSD2, SLC2A4, PHKB, MYPN, PHKA1, SRL, MYLK2, TMOD4, ATP1A2, TXLNB, STAC3, ASB15, NMRK2, MB* |

**Table 5:**
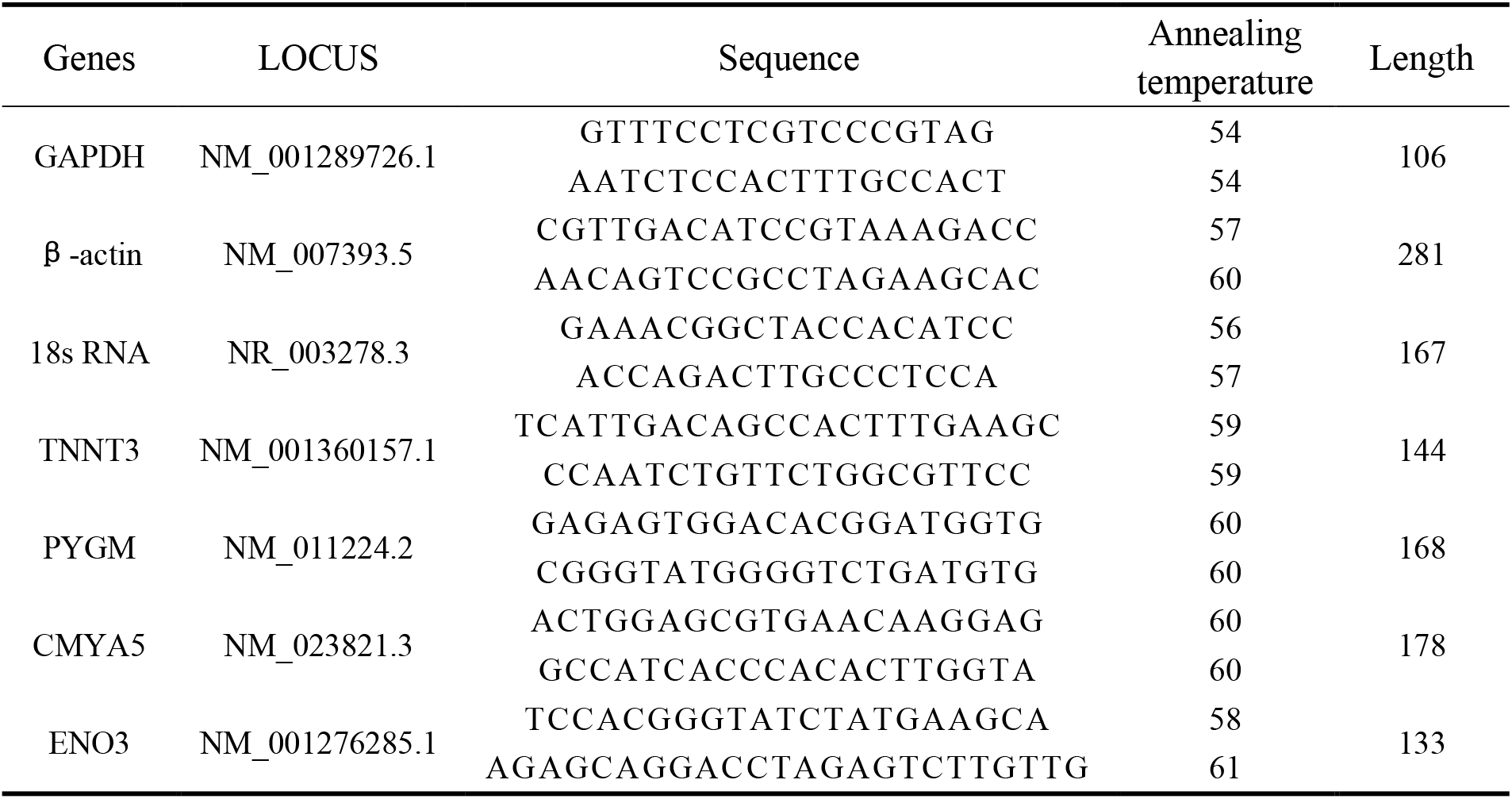
Primer sequences and annealing temperatures used for quantitative PCR.

**Figure 5.**
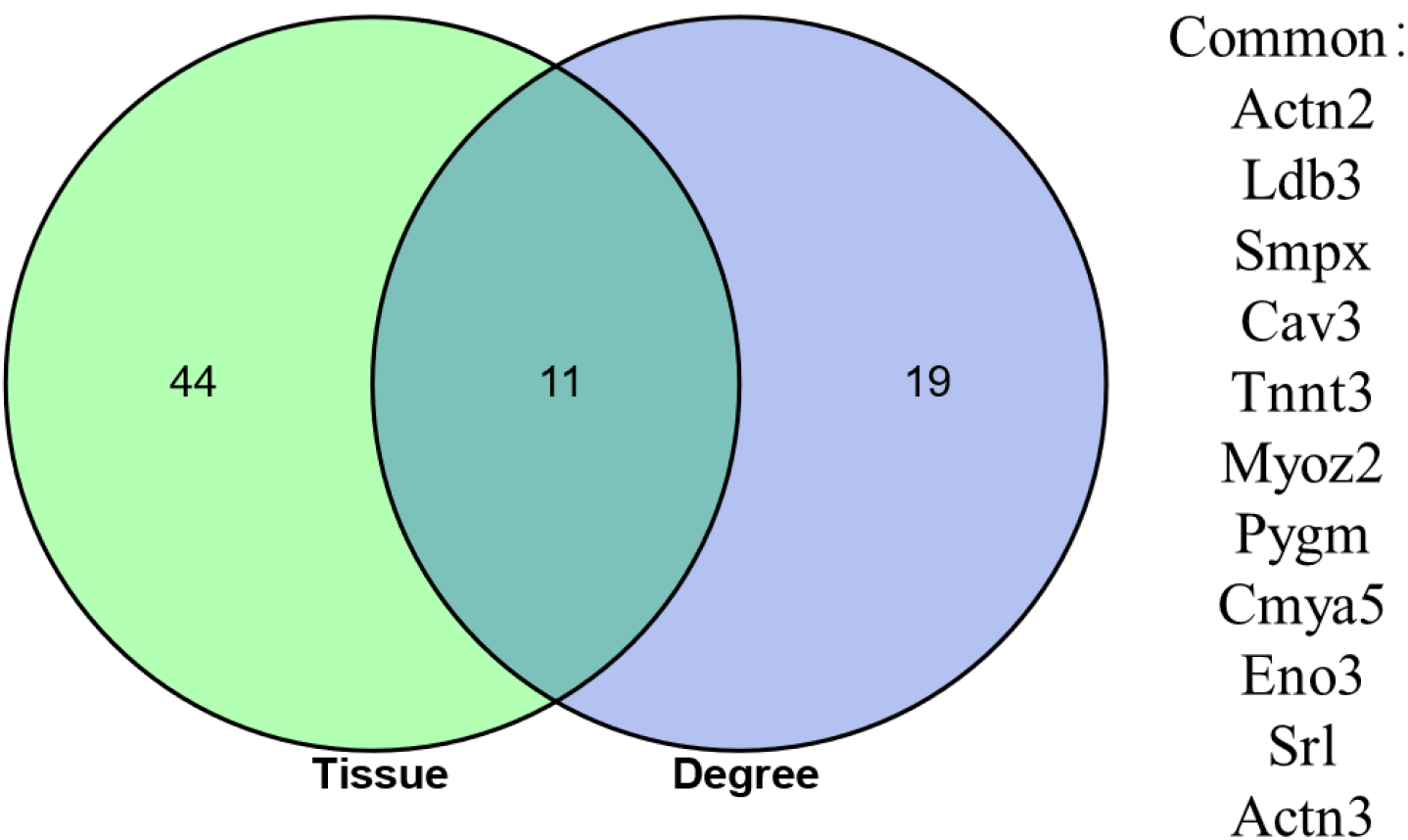
Venn diagram showing the overlap between 55 DEGs identified as having a skeletal muscle specific annotation in DAVID analysis, and the top 30 DEGs identified has having the highest number of connections (degrees) in the PPI network constructed using the online STRING database.

qRT-PCR was performed to determine mRNA expression levels in seven different murine tissues including the heart, liver, spleen, lung, kidney, muscle and adipose tissue to validate muscle specific expression of selected genes. Results from qRT-PCR (Figure 6), confirmed high levels of expression of *TNNT3, PYGM, ENO3, CMYA5* in muscles, reaffirming the validity of the findings in this study. We used *18S rRNA, β-actine* and *GAPDH* as housekeeping genes for the mRNA expression analysis of DEGs in the target tissues. Although *GAPDH* is not considered a very suitable option for using as a reference gene (Li et al., 2017), however, we used triple reference genes for the expression of mRNA levels in all target tissues.

**Figure 6.**
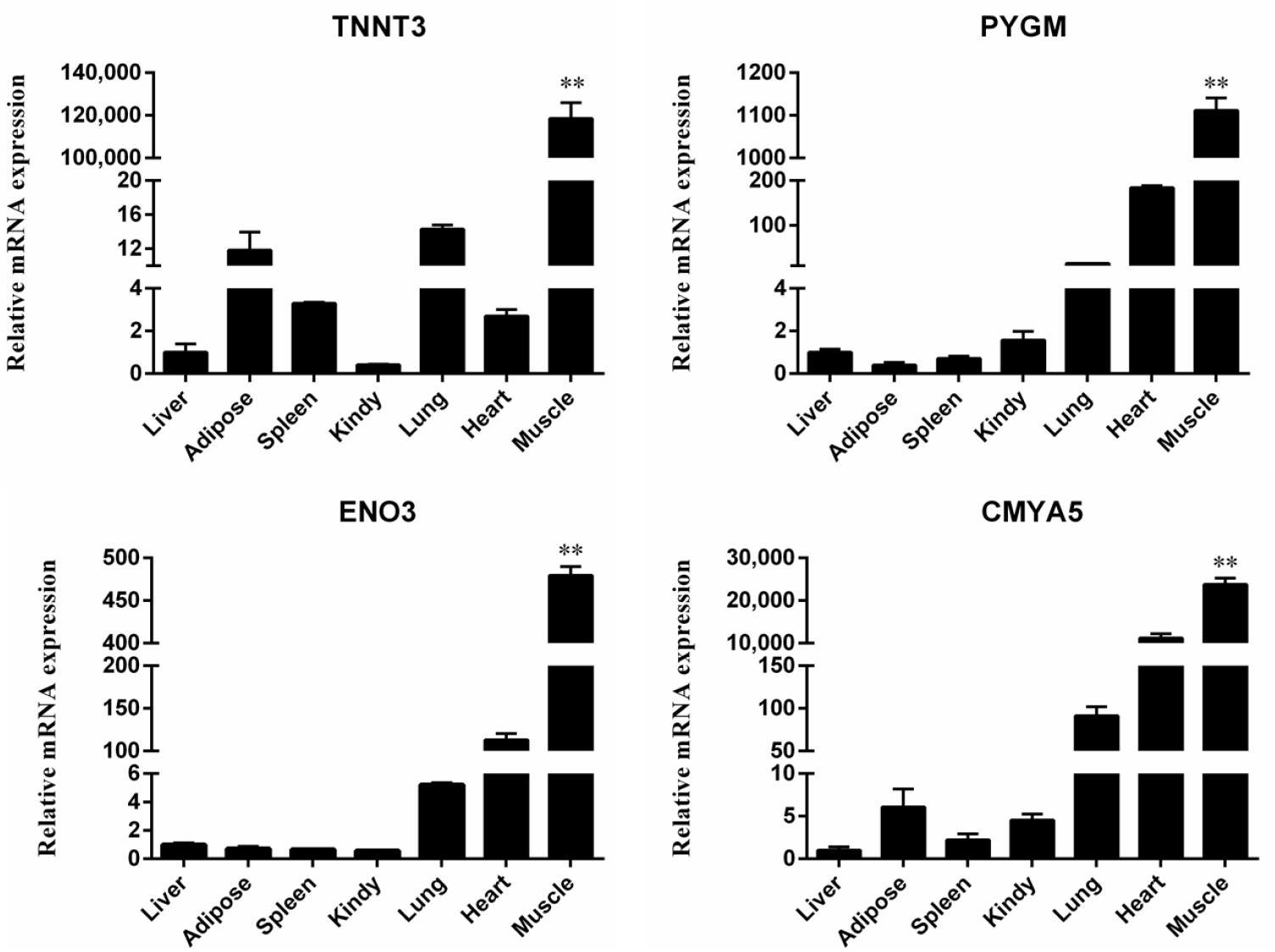
Quantitative real-time PCR validation of differential gene expression. Relative expression levels of four genes (*TNNT3, PYGM, ENO3* and *CMYA5*) in seven different tissue samples. Each bar represents the mean ± SD of the three replicates, and ** represents statistical significance at p < 0.01

## 4. Discussion

The overall aim of the current study was to identify genes specifically expressed in skeletal muscles via bioinformatic analyses of publicly available microarray data, followed by qRT-PCR to validate muscle specific expression of selected genes in an independent set of samples. Bioinformatic analysis of publicly available microarray data resulted in the identification of 272 DEGs with at least a 4-fold expression level relative to expression in 17 other mice tissues. The majority of these genes were upregulated (n=260), and a very small proportion of genes were downregulated (n=12). These results suggest that upregulation of key genes is more crucial for muscle physiology and development, relative to downregulation of specific genes. A previous study (Song et al., 2013) which describes gene expression profiles of different tissues (kidney, liver, lung, heart, muscle, and adipose tissue), also reported that key genes (e.g. *Myot, Tnnc2, Tnni2, Tnnt3, Actn3, Mybpc1, Mybpc2, Myoz1*) are highly upregulated in both human and murine muscles. In our study, we have also found almost all of the above genes to also be highly upregulated (Table 2) in muscles, and therefore our results align with these previous findings. A number of these DEGs have also previously been reported to be involved in muscle development. Some of these genes include *Tnnt3* (Gollapudi et al., 2013; Zhang et al., 2016), *Myh1, Myh2, Myh4* (Ahn et al., 2018; Gaglianone et al., 2020; Wollersheim et al., 2019) and *Actn3* (Miyamoto et al., 2018), *Pvalb*, (Murphy et al., 2012) *Ckmt2*, (Kazak and Cohen, 2020) *Cox8b(Flynn et al*., *2010)*. However, several other genes that were identified to be differentially expressed in muscles have not yet been reported to have a role in muscle development or physiology.

Enrichment and pathway analyses performed identified several GO annotations terms (n=752) to be significantly enriched in the 272 genes identified to be differentially expressed in skeletal muscles. The GO enrichment demonstrated significant involvement of ontologies relevant to muscle development, physiology and function; which in turn accords with findings from DEG analyses. Pathway analysis also identified 29 KEGG pathways, several of which were relevant to muscle development. However, one of the more interesting findings here was that the top four pathways identified were all associated with cardiac muscle physiology and pathology. This could suggest that some genes specifically expressed in skeletal muscles, could be involved in cardiac myopathies.

While the skeletal muscle samples used in this study were derived from 10-12 week old mice, these samples include both satellite cells and skeletal muscle-derived stem cells, which together form the pool of cells required for myogenesis (Mauro, 1961). When satellite cells are activated (e.g. due to muscle injury), they get induced to undergo myogenic differentiation, which in turn requires highly specific temporal and spatial expression patters of different transcription factors and proteins (Tajbakhsh, 2009) that is consistent with findings in this study. Similarly, skeletal muscle stem cell proliferation and muscle differentiation can also be triggered in adults under the influence of hormones like IGF1 (Matsakas and Patel, 2009), which in turn activates a number of downstream pathways including MAPK, PI3K-AKt-mTOR-P70S60K and PI3K-AKt-mTOR-GSKβ signaling pathways (Sandri et al., 2004). Therefore, the identification of several genes, ontologies and pathways associated with muscle development is not surprising.

To affirm the findings from differentially expressed genes DEGs, enrichment and pathway analysis, we constructed a PPI protein interaction network map, consisting of 2,813 edges (interactions) between 247 nodes (proteins). The PPI network map identified several structural proteins and enzymes as core nodules (e.g. *TMOD4, MYL1, MYBPC2, ATP2A1*). When degree scores were computed via Cytoscape network analysis, the top nodes were also mainly comprised of structural genes, and genes involved in muscle physiology and function (e.g. *TNN, ACTN2, LDB3, CKMT2*). Network module analysis via the MCODE plug-in also identified 7 interaction modules. The largest of these modules was comprised of a total of 43 nodes and 775 edges, and included several structural myosin-related (Myh7, Myl2, Myl3, Myh2, Myh4, Myh1) and actin-related (Acta1, Actc1, Actn2, Actn3) proteins. Overall, findings from enrichment, pathway and network analysis were in accord and reaffirmed the involvement of identified DEGs in muscle structure and physiology.

Module analysis of the PPI network identified several genes that have been previously reported to be involved in muscle development (e.g. Module 1, Figure 4C). However, several interesting candidates, whose roles in muscle development are yet to be characterized, were also identified. Examples of such genes include Tripartite motif-containing 54 (*Trim54*), Creatine kinase, mitochondrial 2 (*Ckmt2*), cardiac disease associated 5 (*Cmya5*) and Leiomodin 2 (*Lmod2*). Future research aimed at characterizing the function of these genes could offer novel insights into mechanistic aspects of muscle development and associated pathophysiology.

Finally, we used qRT-PCR to validate the expression patterns of selected genes that were were identified as specifically expressed in skeletal muscles (via DAVID analysis), and were also identified within the top 30 genes of the PPI network (i.e. those having the highest scores). The obtained results are consistent with the results from microarray DEG analyses, which reaffirms the findings from bioinformatic analyses of the microarray data.

## 5. Conclusion

In conclusion, 272 genes with muscle-specific expression profiles were identified in this study, which included several genes widely known to be involved in muscle development and function. Downstream enrichment and pathway analysis identified several muscle specific ontologies and pathways reaffirming findings of differentially expressed genes DEG analysis. Validation of results in an independent set of samples via qRT-PCR also reaffirmed muscle specific expression of selected DEGs. Several of the 272 differentially expressed genes DEGs identified in this study are yet to be functionally characterized in context of muscle development and physiology. Once characterized, these candidate genes could offer new targets for development of mutant mouse models of human muscle associated diseases and disorders. Therefore, future research aimed at investigating the role of these candidate genes in the context of muscle development and physiology is warranted.

## Acknowledgements

All the authors acknowledge and thank their respective institutes & universities.

## Availability of data

All datasets used in this article are public and sources cited accordingly. The screening data respectively Source is available at https://david.ncifcrf.gov https://david.ncifcrf.gov/home.jsp http://kobas.cbi.pku.edu.cn/ http://ggplot2.tidyverse.org https://www.r-project.org/.http://www.bioconductor.org/packages/3.5/bioc/html/limma.html

## Supplementary Materials

The following Supplementary Materials are available at Table S1. Muscle and 17 tissues, S2. GO (Gene Ontology), S3. Enrichment of KEGG pathway, S4. degree Venn diagram in PPI network

